# Seminal plasma exosomes evoke calcium signals via the CatSper channel to regulate human sperm function

**DOI:** 10.1101/2020.05.21.094433

**Authors:** Xiaoning Zhang, Dandan Song, Hang Kang, Wenwen Zhou, Houyang Chen, Xuhui Zeng

## Abstract

Seminal plasma exosomes (SPE) have been proposed to regulate intracellular calcium concentration ([Ca^2+^]i) and sperm function. However, neither the underlying mechanisms by which [Ca^2+^]i is regulated by SPE nor the physiological and pathological significance of the SPE-evoked calcium signal are fully understood. Here, we successfully isolated and characterized SPE by several methods, including transmission electron microscopy, nanoparticle tracking and nanoflow cytometry analysis. Application of SPE dose-dependently increased human sperm [Ca^2+^]i via extracellular Ca^2+^ influx. The Ca^2+^ influx was mediated by the sperm-specific CatSper channel, because the SPE-elevated [Ca^2+^]i was suppressed by a CatSper inhibitor, and SPE potentiated the CatSper current in human sperm. The role of CatSper in the SPE-induced elevation of [Ca^2+^]i was further confirmed by the absence of the SPE-induced [Ca^2+^]i increase and CatSper current in a CatSper-deficient sample. Furthermore, both protein and no-protein components in SPE were shown to contribute to the elevated [Ca^2+^]i, as well as the hyperactivated motility of human sperm. Interestingly, when sperm were stimulated with exosomes derived from asthenozoospermic semen, the elevation of [Ca^2+^]i was significantly lower than that by exosomes isolated from normal seminal plasma. The SPE from normal seminal plasma improved the motility of sperm from asthenozoospermic samples. Taken together, these findings demonstrated that SPE modulates Ca^2+^ signaling and human sperm function by activating a CatSper channel. The application of SPE to enhance sperm motility may provide a new clinical avenue for asthenozoospermic men.

## Introduction

Exosomes are shed from most cell types and exist in a wide variety of body fluids. Increasing evidence indicates that exosomes are important mediators of intercellular communication, rather than non-specific waste from cells [1]. Exosomes can be taken up via specific membrane surface receptors or can release certain signal molecules to regulate recipient cell functions [2, 3]. Mammalian seminal plasma contains membranous exosomes (extracellular vesicles) mainly produced by the epididymis and prostate, traditionally termed epididymosomes or prostasomes, which contain proteins, nucleic acids, and a high content of cholesterol and sphingomyelin [4–7]. Previous studies have demonstrated that seminal plasma exosomes (SPE) and their cargos have antibacterial, antioxidant, and immunosuppressive properties, and may be involved in several biological processes indirectly influencing sperm function [8, 9]. For example, exosomes reduced ROS overproduction and attenuated oxidative stress in sperm [10], and transferred Ca^2+^ signaling factors to regulate progesterone-induced sperm motility [11]. Directly interacting with sperm, human SPE may play pivotal roles in sperm maturation in the epididymis, as well as sperm function and fertilization [12–15]. Interestingly, SPE isolated from oligoasthenozoospermia donors exhibited different miRNA and protein profiles than SPE from normozoospermic exosomes [16, 17]. Furthermore, normal human exosomes promoted sperm motility and capacitation, while this effect was impaired in exosomes derived from donors with asthenospermia [18]. These studies indicate that SPE act as functional regulators of male fertility, and also suggest that dysfunction of exosomes may contribute to male infertility.

Intracellular calcium homeostasis is a key component in the control of sperm motility, and Ca^2+^ influx is important for the maintenance of progressive motility and sperm function [19, 20], especially for the acrosome reaction and hyperactivation [21, 22]. All these processes are indispensable for sperm fertility. Consistent with roles in the regulation of sperm function, SPE increased intracellular Ca^2+^ concentrations ([Ca^2+^]i), thought to be mediated by membrane fusion with sperm rather than ion exchanges and ATP-dependent ion pumps [23, 24]. Only recently has not convincing data been documented concerning the precise mechanism of exosome actions in the sperm [Ca^2+^]i response.

Cation channel of sperm (CatSper), a unique cation channel protein family exclusively expressed in sperm, is composed of at least nine subunits and control Ca^2+^ influx to regulate sperm functions, such as capacitation, chemotaxis, the acrosome reaction and hyperactivation [25]. CatSper channel is essential for male fertility because CatSper-deficient male mice and men with mutations in CatSper genes exhibit infertility due to lack of sperm hyperactivation [26]. CatSper can be activated by many physiological substances, such as progesterone, prostaglandins and certain chemicals [27, 28]. Thus it has been considered as a polymodal chemosensor that regulates sperm function and fertility [28]. However, the role of CatSper in exosome-induced [Ca^2+^]i remains unknown. Also, investigation is required to determine the components of exosomes that exert the regulatory function on human sperm function.

The present study investigated the mechanism by which SPE regulate Ca^2+^ signaling in human sperm, with a focus on the involvement of CatSper. In addition, the functional and pathological significance of SPE-evoked [Ca^2+^]i responses was also examined, to provide insight into the potential clinical application of SPE.

## Materials and Methods

### Reagents

Human tubal fluid (HTF) medium were purchased from Nanjing Aibei Biotechnology Co., Ltd (Nanjing, China). Fluo-4 AM and Pluronic F-127 were obtained from Molecular Probes (Eugen, OR, USA). Triton X-100 and Proteinase K (Pro K) were obtained from Solarbio (Beijing, China). Mibefradil (Mi), progesterone (P4), 1, 2-bis (o-aminophenoxy) ethane-N, N, N’, N’-tetraacetic acid (BAPTA), EGTA and A23187 were purchased from Sigma-Aldrich (Shanghai, China). CD9 and CD63 antibodies were obtained from proteintech (Wuhan, China). Prostaglandin E1 (PGE1) and E2 (PGE2) were obtained from Aladdin (Shanghai, China) and TCI Development Co., Ltd (Shanghai, China), respectively.

### Sperm sample preparation

The collection of semen samples and experiments in this study were approved by the Institutional Ethics Committee on human subjects of the Nanchang Reproductive Hospital and Jiangxi Maternal and Child Health Hospital. Informed consent documents were signed by the donors. Semen viscosity and volume, sperm motility, and sperm concentration were assessed according to the WHO laboratory manual for the examination and processing of human semen (WHO 2010, https://www.who.int/reproductivehealth/publications/infertility/9789241547789/en/).

Samples were subjected to high-saline solution (HS, 135 mM NaCl, 5 mM KCl, 1 mM MgSO4, 2 mM CaCl2, 20 mM HEPES, 5 mM glucose, 10 mM lactic acid and 1 mM Na-pyruvate, adjusted to pH 7.4 with NaOH) for washing or a swim-up purification method for the experiments, as described in previous studies with some modifications [29, 30].

### Exosome extraction

After semen was centrifuged at 7,850 g for 30 min at 4°C, the supernatant was collected and stored at −80°C until use. Before isolation, seminal plasma was filtered successively with 0.45 and 0.22 μm MF-Millipore™ membranes (Millipore, MA, USA). The filtered seminal plasma was centrifuged at 10,000 g for 30 min at 4°C to remove cell debris and other impurities. Remaining supernate was centrifuged at 68,000 g for 90 min at 4°C. Then, the precipitate was suspended in 35 mL of 0.32 M sucrose and centrifuged at 100,000 g for 90 min at 4°C. Next, the precipitate was washed with PBS at 100,000 g for 90 min at 4°C. Finally, the exosomal pellets were resuspended in 3.5 mL HS and stored at −80°C for further study.

### Nanoflow analysis

The diameter range and concentration of exosomes were analyzed with a Flow NanoAnalyzer N30 (NanoFCM Inc, Xiamen, China). Marker protein expression in exosomes was analyzed by nanofluidic methods. The exosomes were labeled by CD63 antibody (1:100) and corresponding secondary antibody (1:5000, Invitrogen, CA, USA). During the experiment, primary or secondary antibodies alone were added to samples as controls. In addition, exosomes with secondary antibody alone, and CD63 antibody with PBS were used as background controls. After a final dilution (1:5000), each sample was analyzed by a Flow NanoAnalyzer N30 (NanoFCM Inc, Xiamen, China).

### Transmission electron microscopy (TEM)

The exosomes were observed with a TEM (JEM-1200EX, Japan Electronics Co., Ltd.). First, 5–10 μL exosomes were sedimented on copper mesh for 3 min and liquid absorbed with filter paper. After the sample was rinsed with PBS, it was dyed with phosphotungstic acid. Finally, it was dried at room temperature for 2 min and images were captured on the TEM (electron microscopy operating voltage 80–120 kV).

### Computer-aided sperm analysis system (CASA)

To evaluate the effect of exosomes on sperm motility, seminal plasma and exosome-free seminal plasma were prepared. Seminal plasma was prepared as follows: cell debris and sperm were removed from intact seminal plasma by centrifugation and > 220 nm vesicles were filtered with a 0.22 μm MF-Millipore™ membrane (Millipore, MA, USA). The filtered seminal plasma was centrifuged at 68,000 g for 90 min and regarded as exosome-free seminal plasma. Sperm was resuspended in seminal plasma and exosome-free seminal plasma, and then placed in a 5% carbon dioxide incubator (SANYO Electric Co., Ltd., Osaka, Japan) at 37°C for different time periods. Sperm parameters were measured by CASA (WLJY-9000, WeiLi. Co., Ltd., Beijing, China). Total motility, progressive motility (PR), curvilinear velocity (VCL), straight-line (rectilinear) velocity (VSL), average path velocity (VAP), linearity (LIN) and other sperm parameters were recorded.

### Evaluation of the ability of sperm to penetrate viscous media

The environment of the female reproductive tract was simulated with 1% methylcellulose, because sperm complete fertilization with an egg after migration through the female reproductive tract. After sperm were selected by a swim-up purification procedure (30 min, 37°C) the sperm in the upper fractions were washed twice in HS buffer and adjusted to a concentration of 2.0 × 10^7^ cells/mL. Exosomes or other drugs were added after capacitation for 2 h (progesterone was added after 2.5 h). Three h later, 20 μL of 1% methylcellulose was drawn into a glass capillary tube, and the capillary placed in the mixed sperm. The number of sperm in 2 and 3 cm (from the base) of the capillary tube were recorded after 1 h.

### Assessment of sperm hyperactivation (HA)

Sperm were capacitated in HTF medium for 3 h and treated with exosomes and drug for 1 h at 37°C. After capacitation, 8 μL of sperm samples were used to assess motion characteristics via the CASA system (Hamilton Thorne, Inc., Florida, USA) and at least 200 spermatozoa were counted for each assay. Aliquots of 8 μL non-capacitated sperm were used as a non-hyperactivated control. The ALH, LIN, VCL and other parameters were recorded. Hyperactivation was defined as cells with a curvilinear velocity (VCL) ≥ 150 μm/s, linearity (LIN) < 50%, and lateral head displacement (ALH) ≥ 7 μm, as described previously (Alasmari, et al., 2013). The percentage of hyperactivated sperm was calculated as the number of hyperactivated sperm/all motile sperm.

### Assessment of intracellular calcium

Intracellular calcium levels in sperm were monitored with Fluo-4 AM dye and F127. In brief, the sperm were washed 3 times with HS and incubated with Fluo-4 AM (2 μM) and F127 (0.1%) in HS for 30 min at 37°C in darkness. After washing the cells with HS two times, sperm was exposed to exosomes or drugs and fluorescence intensity was assessed by the EnSpire^®^ Multimode Plate Reader (PerkinElmer, Waltham, MA, USA) with excitation and emission wavelengths of 488 and 525 nm, respectively. To determine a change in the calcium signal, the sperm suspension was added to a 96-well plate, then the base fluorescence intensity (F0) was measured. After the drug was added, the fluorescence (F) was measured, and the change in sperm [Ca^2+^]i was calculated by ΔF/F0 (%) = (F - F0)/F0 × 100%.

### Single cell calcium imaging technology

The samples were purified by a 50% Percoll solution (Percoll:HSA:EBSS:ddH20 = 5:1:1:3). After centrifugation at 500 g for 15 min, the sperm was collected, washed once with HS and resuspended in HS. The sperm was incubated with Fluo-4 AM and F127 in a 5% carbon dioxide incubator at 37°C for 30 min. After excess dye was washed off with HS, the sperm was adjusted to a concentration of 2.0 × 10^7^ cells/mL.

Then 200 μL of sperm suspension was placed evenly into a dish bottom. The sample was adhered the wall and kept away from light for 30 min. After excess fluid was poured off, sperm in the dish were washed with HS. Finally, [Ca^2+^]i was recorded in a single-cell calcium imaging system (OLYMPUS, Tokyo, Japan) before and after exosome treatment.

### Sperm patch-clamp recordings

The whole-cell patch-clamp technique was applied to record human sperm CatSper currents as previously described (Zeng et al., 2011). Seals were formed in the sperm cytoplasmic droplet or the neck region using a 15–30 MΩ pipette. The pipette solution for recording CatSper currents contained 135 mM Cs-Mes, 10 mM Hepes, 10 mM EGTA, and 5 mM CsCl, adjusted to pH 7.2 with CsOH. Then, the transition into whole-cell mode involved the application of short (1 ms) voltage pulses (400–650 mV) combined with light suction. The currents were stimulated by 1 s voltage ramps from −100 to +100 mV from a holding potential of 0 mV. For recording the monovalent current of CatSper, divalent-free (DVF) solution (150 mM NaCl, 20 mM HEPES, and 5 mM EDTA, pH 7.4) was used to record basal CatSper monovalent currents. Then, different concentrations of exosomes and progesterone were perfused to record CatSper currents. The currents were analyzed with a Clampfit (Axon, Gilze, Netherlands) and figures were plotted with Grapher 8 software (Golden Software, Inc., Golden, Colorado).

### Western blotting

The extracted exosomes were added to 5 × protein loading buffer and boiled at 95°C for 5 min to denature proteins for later use. Each protein sample was subjected to SDS-PAGE and transferred to a PVDF membrane, which was then blocked with 3% BSA for 1 h. Next, the membrane was incubated with CD63 and CD9 antibody (1:1000) at 4°C overnight, then washed three times with TBST. The secondary antibody was added and incubated for 1 h. Then the PVDF membrane was washed three times with TBST for 5 min, then exposed to ECL chemical reagent and examined by a gel imaging system (Bio-Rad, CA, USA).

### ELISA

The content of prostaglandins in exosomes was detected with PGE1 and PGE2 enzyme-linked immunosorbent assay kits from Wuhan Shenke Experimental Technology Co., Ltd. (Wuhan, China) and Nanjing Jiancheng Bioengineering Institute (Nanjing, China), respectively, according to the operating instructions.

### Statistical analysis

Data are expressed as the mean ± SEM. Differences between the controls and treated samples were assessed with the unpaired Student’s t test using the statistical software GraphPad Prism (version 5.01, GraphPad Software, San Diego, CA, USA). P < 0.05 was regarded as statistically significant.

## Results

### Identification of exosomes from human semen

The SPE were isolated from 12–15 different donors by ultracentrifugation and had an average diameter of 94.9 ± 21.8 nm (Figure 1(a)). The SPE size ranged from 50 to 200 nm, with most < 100 nm and more than 4.67 × 10^13^ particles/mL were examined by the Flow NanoAnalyzer N30 (Figure 1(b)). After treatment with Triton-X 100, particle numbers decreased by about 18-fold to 2.66 × 10^12^ particles/mL and many particles became smaller (Figure 1(c)). The Flow NanoAnalyzer and western blotting analysis showed the presence of universal exosome markers, CD9 and CD63, in the isolated exosomes (Figure 1(d)–1(f)). TEM further confirmed that the exosomes contained a single lipid bilayer and appeared as round particles in the expected size range of exosomes (Figure 1(g)).

**Figure 1.**
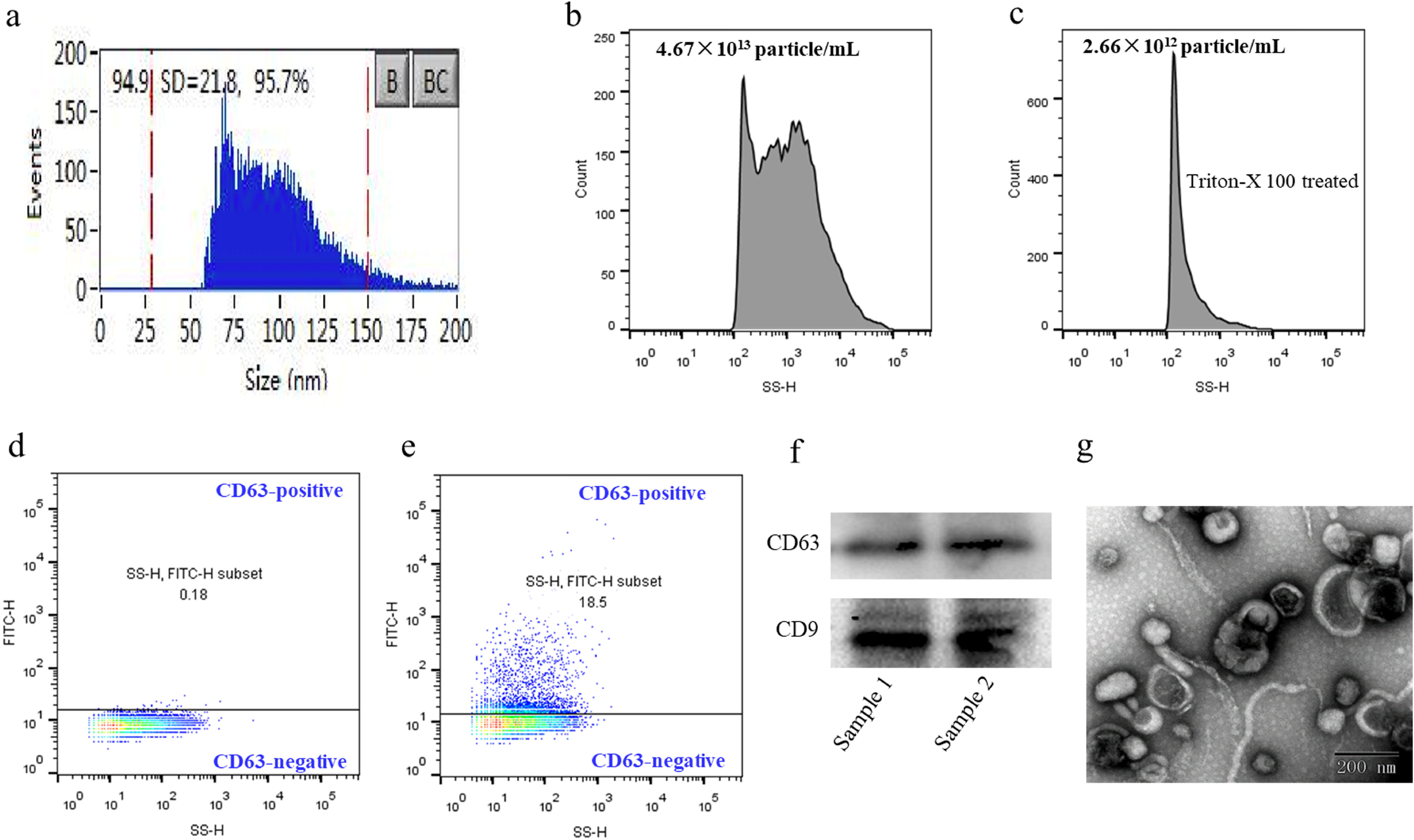
Characteristic of SPE. (a) The diameter range of exosomes. After ultracentrifugation, the exosomes were analyzed with Flow NanoAnalyzer. (b) (c) The characteristic and concentration of particles. Intact exosomes and triton X-100-treated (1%, 30 min) exosomes were detected by the Flow NanoAnalyzer. (d) (e) Expression of CD63 in SPE. Exosomes were subjected to immunofluorescence staining with antibodies directed against CD63 and then measured with Flow NanoAnalyzer. (f) Expression of CD63 and CD9, exosome markers, in SPE. Exosomes from two different donors were subjected to SDS-PAGE and immunoblot analysis with an antibody directed against CD63 and CD9. Western blots are representative of N = 3 experiments. (g) Morphology of exosomes by transmission electron microscope. It showed that there is a typical feature of exosomes with intact lipid bilayers and saucer-like structure of particles.

### Exosomes elevated [Ca^2+^]i signals via extracellular Ca^2+^ influx

It was reported that human spermatozoa [Ca^2+^]i increased after fusion of sperm with exosomes [31], however, the source of the [Ca^2+^]i burst remained to be clarified. The present study investigated the effect of isolated exosomes on human sperm [Ca^2+^]i. The results indicated that exosomes increased human sperm [Ca^2+^]i within 1 min in a dose-dependent manner (Figure 2(a) and 2(b)). The concentration of applied exosomes was close to physiological conditions when the ratio of exosome:sperm was 1:1. Next, we further confirmed the exosome-evoked Ca^2+^ response by single-sperm [Ca^2+^]i imaging technology (Figure 2(c)). To determine whether the exosome-induced increase in [Ca^2+^]i was due to Ca^2+^ influx or the mobilization of Ca^2+^ stores, we measured the [Ca^2+^]i of sperm exposed to a Ca^2+^-free medium containing Ca^2+^ chelators BAPTA or EGTA. Under these conditions, we found that exosomes had no effect on [Ca^2+^]i (Figure 2(d) and 2(e)), indicating that the exosome-induced human sperm [Ca^2+^]i resulted from extracellular Ca^2+^ influx.

**Figure 2.**
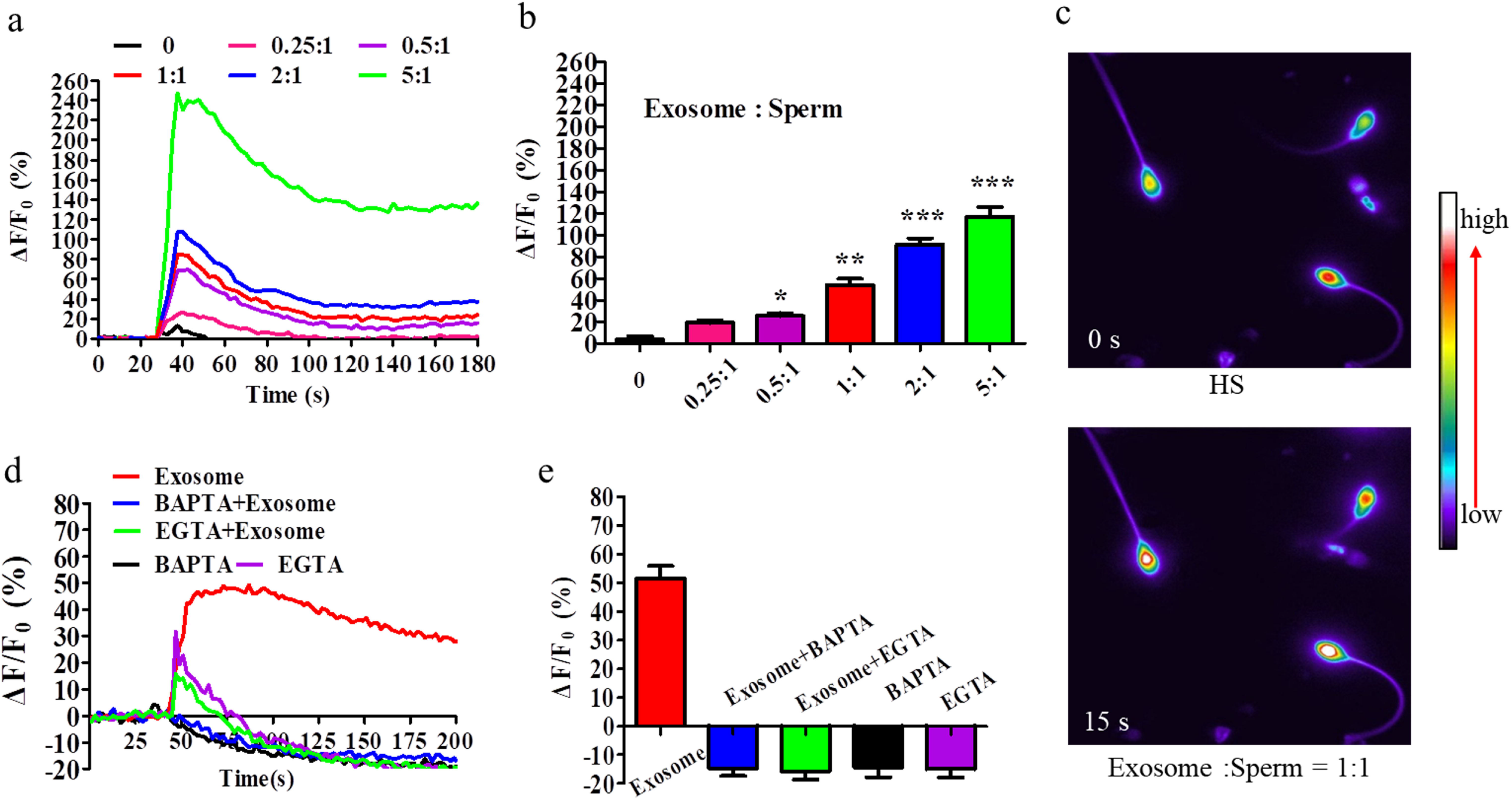
Exosomes induced [Ca^2+^]i increase via extracellular Ca^2+^ influx. (a) Examples of [Ca^2+^]i change was illustrated with the time frame. (b) The statistical analysis of the effects of different concentrations of exosomes isolated by ultracentrifugation on sperm [Ca^2+^]i. Sperm [Ca^2+^]i was monitored after loading cells with Fluo-4 AM-AM (2 μM) and F127 (0.1% w/v) and the fluorescence intensity of the sperm was detected by microplate reader before and after adding the different concentrations of exosomes. (c) Single-sperm [Ca^2+^]i imaging indicated a [Ca^2+^]i response in sperm after exosome treatment. Sperm [Ca^2+^]i was stained with Fluo-4 AM-AM (2 μM) and F127 (0.1% w/v), and the fluorescence intensity of the sperm was visualized and detected before and after adding exosomes. (d) Effect of exosomes on sperm [Ca^2+^]i in Ca^2+^-free buffer. Sperm [Ca^2+^]i was determined before and after adding exosomes in the HS buffer with presence or absence of BAPTA (12 mM) or EGTA(5 mM). (e) The statistical analysis of the amplitude ΔF/F0 of [Ca^2+^]i change from (d) time-course traces (n = 3). Data were presented as mean ± SME (n = 10). *P < 0.05, **P < 0.01, ***P < 0.001 versus HS group.

### CatSper-mediated exosome-induced [Ca^2+^]i response

In human spermatozoa, CatSper is the predominant Ca^2+^ channel responsible for Ca^2+^ influx [26]. Hence, we examined whether CatSper was involved in the exosome-induced human sperm [Ca^2+^]i burst via extracellular Ca^2+^ influx. The CatSper blocker, Mi, significantly impaired the exosome-activated Ca^2+^ response, suggesting that CatSper was involved in the exosome-induced increase in sperm [Ca^2+^]i (Figure 3(a) and 3(b)). To further investigate this possibility, the whole-cell patch-clamp technique was applied to examine the activation effect of exosomes on sperm CatSper. The results showed that exosomes potentiated CatSper currents in normal sperm samples (Figure 3(c) and 3(d)), supporting the role of CatSper in the exosome-induced increase of [Ca^2+^]i. To clarify the overall role CatSper in mediating exosome-induced Ca^2+^ influx, we took advantage of an infertile sperm sample lacking the CatSper2 subunit; thus lacking the CatSper current [32]. The data showed that exosomes did not increase [Ca^2+^]i and CatSper current in the CatSper-deficient sample (Figure 3(e) and 3(f)). As a control, a CatSper independent Ca^2+^ signal activator (A23187) was found to increase sperm [Ca^2+^]i (Figure 3(f)). Collectively, these results suggested that CatSper controlled exosome-induced [Ca^2+^]i signal responses.

**Figure 3.**
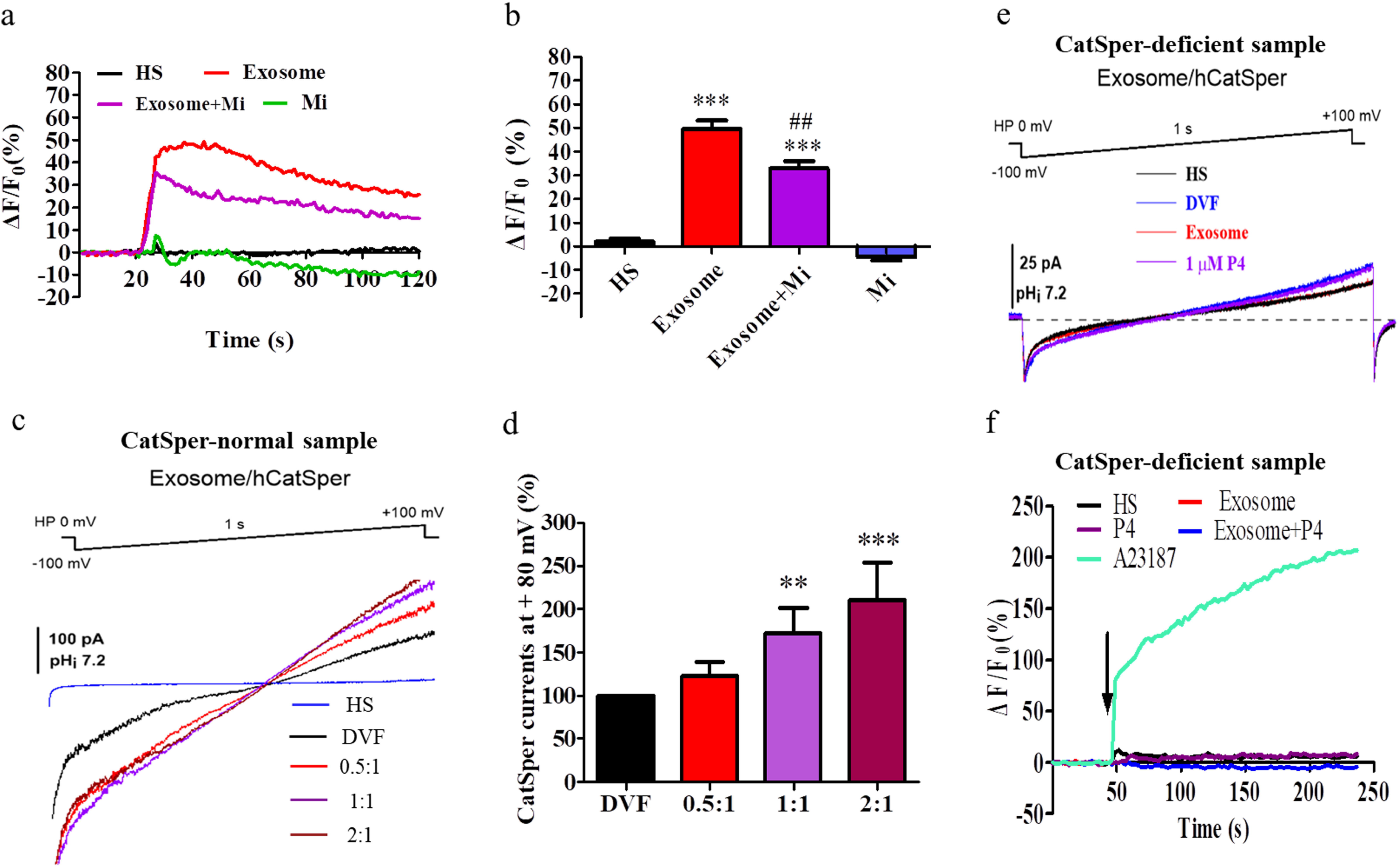
CatSper mediated exosome-induced [Ca^2+^]i signal. (a) The CatSper blocker, Mi, impaired the exosome-activated [Ca^2+^] response. Sperm [Ca^2+^]i was monitored after loading sperm with Fluo-4 AM-AM (2 μM) and F127 (0.1% w/v) and the fluorescence intensity of the sperm was detected by microplate reader after treating sperm with exosomes in the presence or absence of Mi (30 μM). (b) The statistical analysis of the amplitude ΔF/F0 of [Ca^2+^]i change from (a) time-course traces (n = 5). (c) Exosomes activated CatSper current in normal human sperm but not in (e) CatSper-deficient sperm. CatSper currents were examined by whole-cell clamp technique from −100 to +100 mV as described in Materials and Methods Section. Different concentrations of exosomes diluted in DVF (divalent free) buffer was perfused to record exosome-induced CatSper currents. DVF solutions were used to record baseline monovalent CatSper currents. (d) Statistical analysis of the mean CatSper currents at +100 mV (positive) and −100 mV (negative) for (c). (f) [Ca^2+^]i in CatSper-deficient sperm have no response to exosomes. After staining sperm with Fluo-4 AM-AM (2 μM) and F127 (0.1% w/v) for 30 min, sperm [Ca^2+^]i was detected before and after adding exosomes (sperm: exosome = 1:1) and/or P4 (2 μM). A23187, a Ca^2+^ signal activator with CatSper independent, (1 mM) served as control. Data were presented as mean ± SME (n = 5). ###P < 0.001 versus exosome group. *P < 0.05, **P < 0.01, ***P < 0.001 versus DVF group.

### Exosome-loading cargos provoked Ca^2+^ signals

To investigate the specific exosomal cargos that controlled calcium signals in human sperm, we obtained supernatants and precipitates from 0.02% triton 100-X-treated exosomes using ultracentrifugation. Only the supernatant components induced the sperm [Ca^2+^]i signal (Figure 4(a) and 4(b)). Furthermore, ultrasonic-disrupted exosomes had a stronger stimulatory effect inducing the [Ca^2+^]i signal compared with intact exosomes (Figure 4(c) and 4(d)). To investigate whether the exosome-bearing proteins mediated the Ca^2+^ effects, the supernatant from exosomes treated with pro K at 58°C overnight (18 h) reduced the sperm [Ca^2+^]i signal compared with the unheated supernatant component (Figure 4(e) and 4(f)). Furthermore, high temperature denaturation of exosome supernatant also reduced the elevation of [Ca^2+^]i (Figure 4(g) and 4(h)). There were also no Ca^2+^ responses to pro K-treated exosomes and exosome supernatant in a Ca^2+^-free medium (Figure S1). These data indicated that both exosome proteins and other non-protein components promoted the [Ca^2+^]i increase via extracellular Ca^2+^ influx. Considering prostaglandins detected in exosomes were reported to increase [Ca^2+^]i signals [32, 33], we investigated the presence of PGE1 and PGE2 in exosomes (Figure S2(a)). We found that both prostaglandins were present in isolated exosomes, although the detected concentrations were lower than the concentrations required to induce a [Ca^2+^]i effect in sperm (Figure S2(b)–(e)). We then examined the role of the intact membrane structure of exosomes for the stability of their prostaglandin cargos. Our data suggested that exosome membranes maintained cargo stability and long-term physiological activity, because triton X-10-treated exosomes had no detectable PGE1 levels and higher PGE2 levels (Figure S2(a). We propose that exosomes provide the necessary cargo stability to evoke a Ca^2+^ signal.

**Figure 4.**
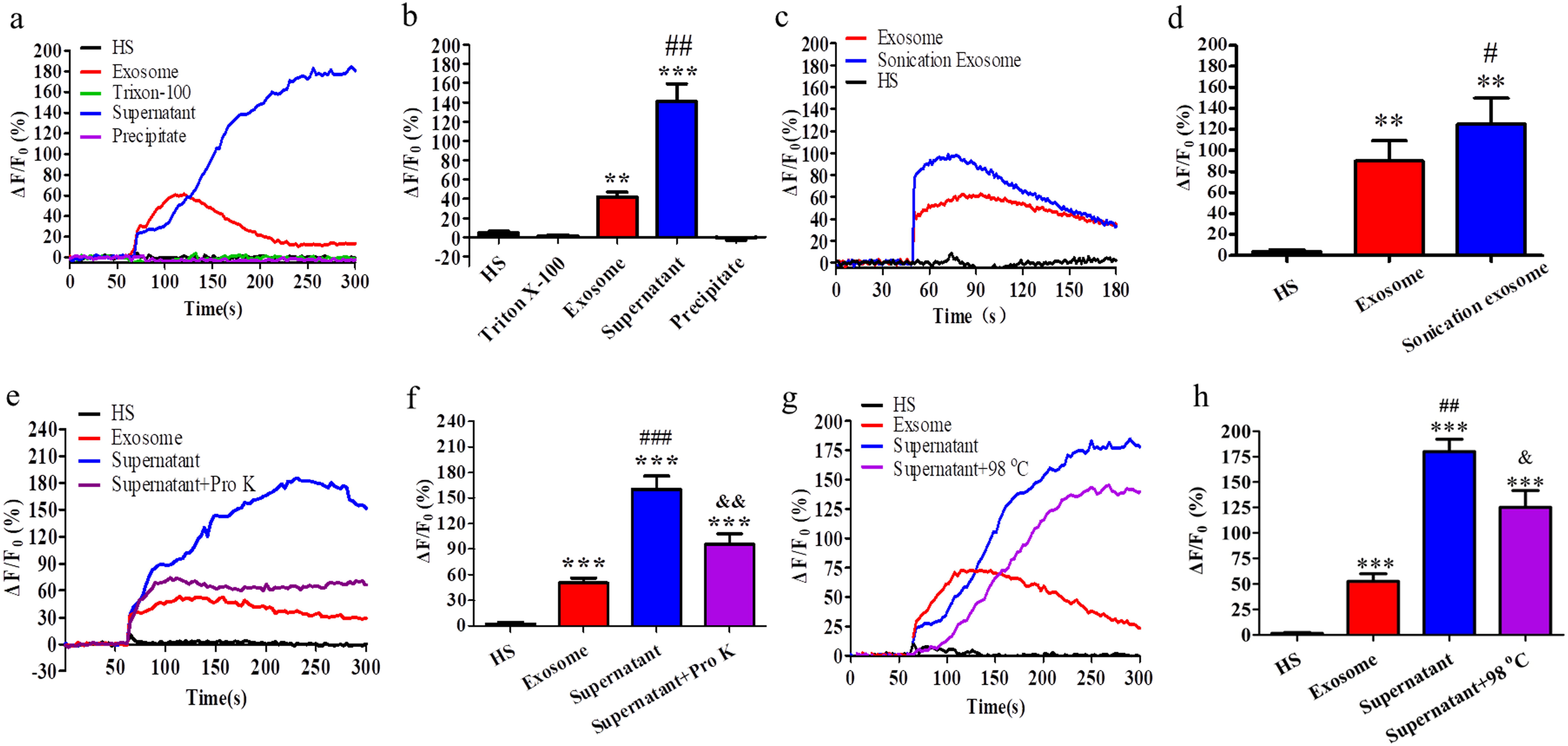
Exosomes-loading protein and non-protein components provoked [Ca^2+^]i signal. (a) Effect of exosome components on sperm [Ca^2+^]i. (c) Effect of ultrasonic on exosome-induced Ca^2+^ level burst. (e) Effect of enzymolysis with Pro K (protease K) on exosome-induced Ca^2+^ level burst. (g) Effect of Pro K on exosome supernatant-induced Ca^2+^ burst. (i) Effect of heating on exosome-induced Ca^2+^ level burst. (k) Effect of long-term conservation on exosome-induced Ca^2+^ level burst. (b) (d) (f) (h) (j) (l) Statistical analysis of the amplitude of the Ca^2+^ changes from (a) (c) (e) (g) (i) (k), respectively. Triton X-100 (0.002%)-penetrated exosomes were ultra-centrifuged to obtain supernatants and precipitates. After stimulating sperm with exosomes, supernatants, precipitates, ultrasonic-, heating- and Pro K-treated exosomes, sperm [Ca^2+^]i was monitored by a microplate reader with Fluo-4 AM-AM (2 μM) staining. Data were presented as mean ± SME (n = 8). #P < 0.05, ##P < 0.01 versus exosome group. *P < 0.05, **P < 0.01, ***P < 0.001 versus HS group.

### Exosomes facilitated sperm hyperactivation

In view of CatSper mainly regulates sperm hyperactivation [34], we examined the effect of the CatSper-mediated increase in exosome-induced [Ca^2+^]i upon sperm hyperactivation. As shown in Figure 5(a), exosomes significantly promoted hyperactivation. In addition, the ability of human spermatozoa to penetrate artificial viscous media, another hyperactivation-related parameter, was enhanced by exosomes and impaired by the CatSper inhibitor Mi (Figure 5(b)). Treatment of exosome supernatant with Pro K or heating significantly impaired the hyperactivated sperm count and the ability of sperm to pass through viscous media (Figure 5(c)–5(f)), implying that exosome-induced sperm hyperactivation results from both exosomal protein and non-protein components.

**Figure 5.**
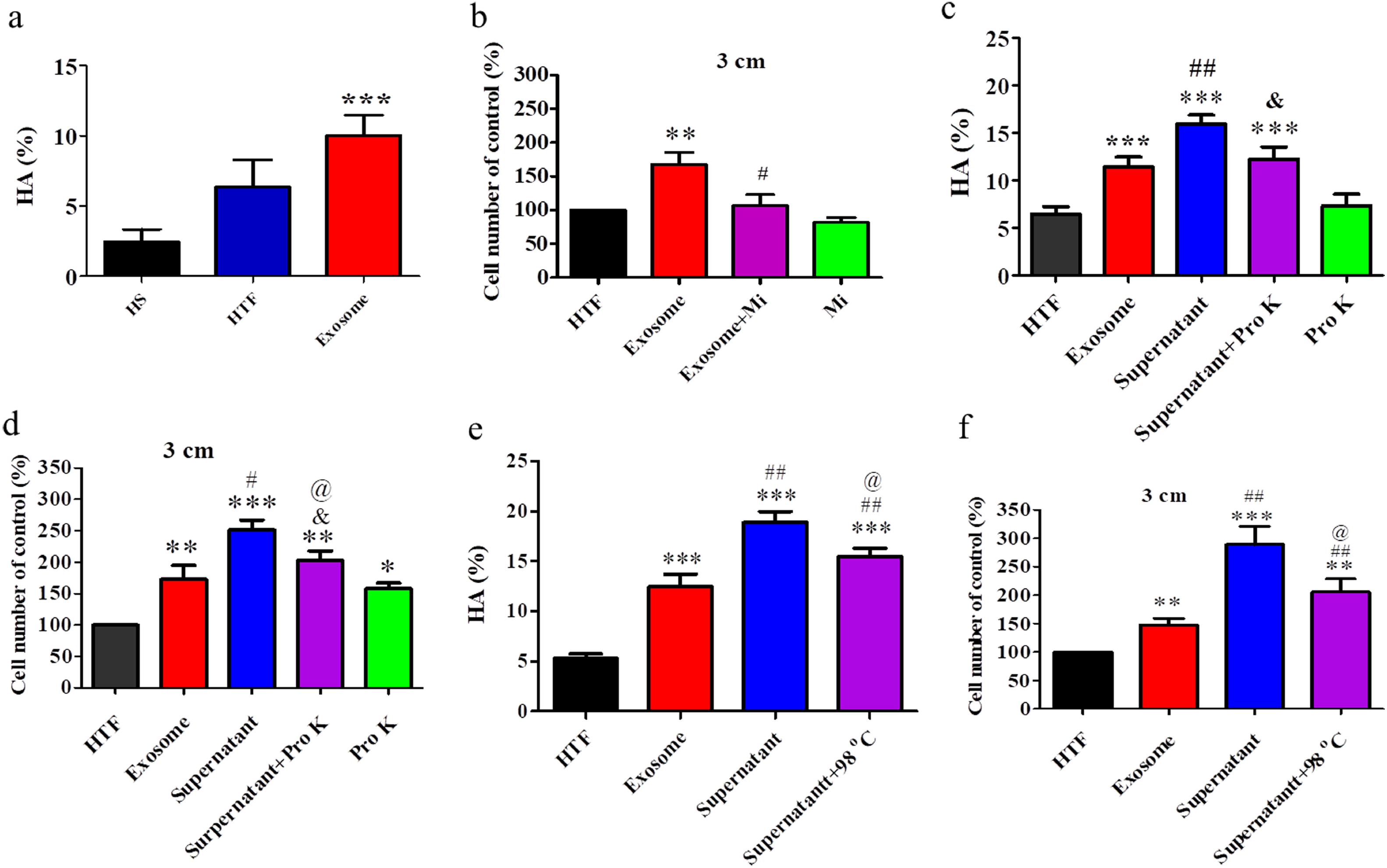
Exosomes and its cargos promoted human sperm hyperactivation. (a) The effect of exosomes on human sperm hyperactivation. (b) Mi impaired exosome-induced the ability of penetration into viscous medium. The exosome supernatant-induced (c) hyperactivation and (d) the ability of sperm to penetrate into artificial viscous media were attenuated by Pro K. The exosome supernatant-induced (e) hyperactivation and (f) the ability of sperm to penetrate into artificial viscous media were attenuated by heating. Hyperactivation was assessed via the CASA system and 200 spermatozoa were counted at least for each assay. The sperm was capacitated in HTF medium for 3 h and treated with exosomes for 1 h at 37°C. Hyperactivation was defined as those cells with a curvilinear velocity (VCL) ≥ 150 μm/s, linearity (LIN) < 50%, and lateral head displacement (ALH) ≥ 7 μm as described in Materials and Methods Section. The ability of sperm to penetrate into artificial viscous media was evaluated as the following protocol. After sperm were selected by a swim-up purification procedure (30 min, 37 °C) and the better vigor sperm in upper fractions were washed twice in HS buffer. Exosomes or treated supernatants were added after capacitation for 2 h. Three h later, 20 μL of 1% methylcellulose was inhaled into the capillary glass tube, and the capillary was placed in the mixed sperm. The number of sperms 3 cm in the capillary from the base of the tube was recorded after 1 h. Data were presented as mean ± SME (n = 5). *P < 0.05, **P < 0.01, ***P < 0.001 versus no treatment group (HTF). ##P < 0.01 versus exosome-treated group.

### Exosomes improved asthenozoospermic sperm motility

Considering exosomes facilitate sperm motility [18] and function via calcium signals, we further investigated a potential clinical application of exosomes. We found that the elevation of [Ca^2+^]i was significantly lower in exosomes derived from asthenozoospermia than from normal seminal plasma (Figure 6(a) and 6(b)). Next, the effects of exosomes upon sperm motility parameters in seminal plasma were investigated by CASA. Sperm total motility, progressive motility, VCL, VSL and VAP parameters were markedly higher in normal seminal plasma than in exosome-free seminal plasma isolated after ejaculation (Figure S3). Consistent with these results, exosomes, especially those derived from normozoospermic semen, improved sperm progressive motility and promoted the penetration of asthenozoospermic sperm into viscous medium (Figure 6(c) and 6(d)). Therefore, exosomes partly rescued sperm motility and sperm function impaired by exosome dysfunction.

**Figure 6.**
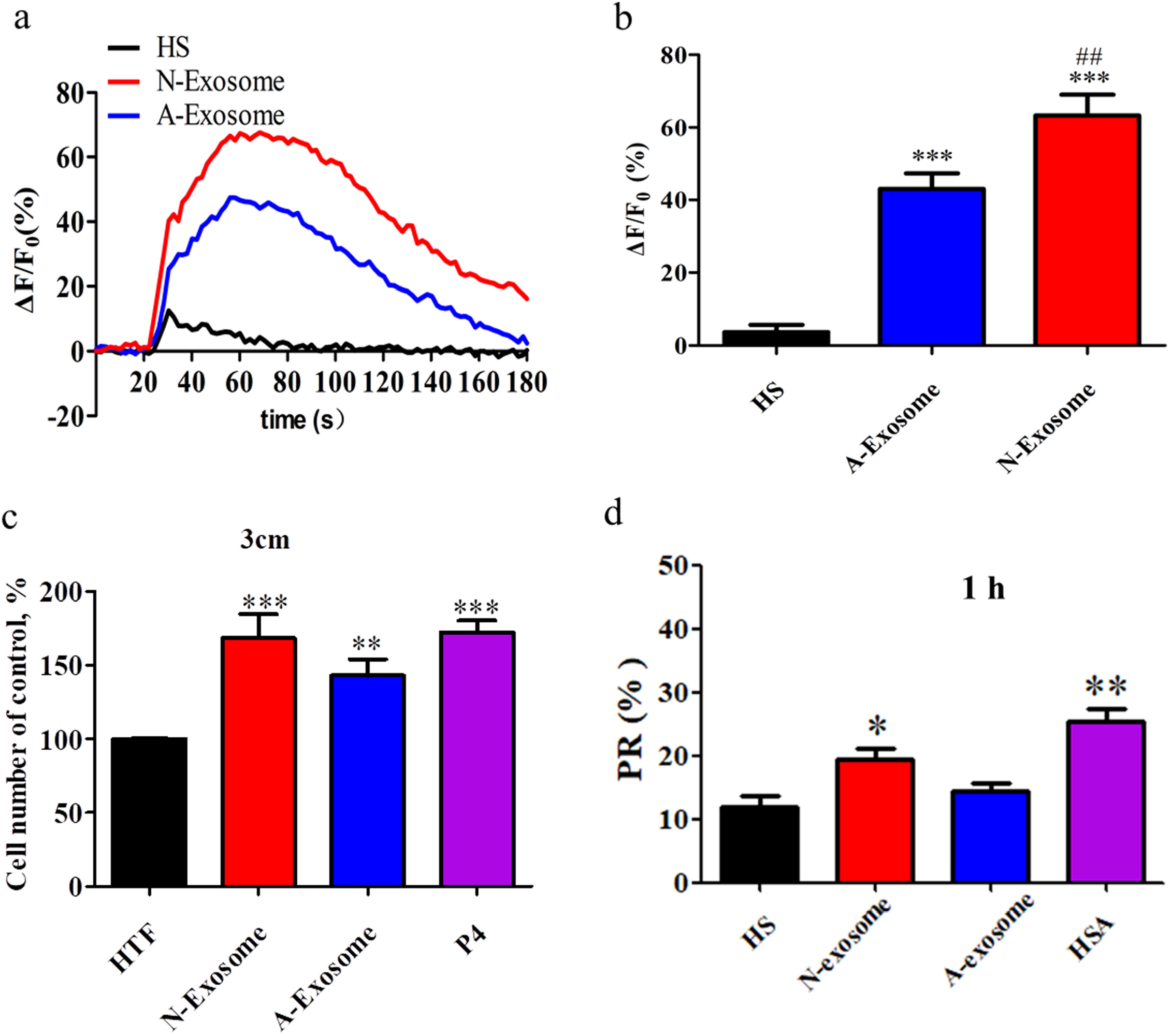
The effects of exosomes on sperm motility and [Ca^2+^]i signal in asthenozoospermia. (a) Effect of exosomes on sperm [Ca^2+^]i. Sperm [Ca^2+^]i was monitored after loading cells with Fluo-4 AM-AM (2 μM) and F127 (0.1% w/v) and the fluorescence intensity of the sperm was detected by microplate reader before and after adding the different concentrations of exosomes extracted from normozoospermic or asthenozoospermic patients. (b) Statistical analysis of the amplitude of the Ca^2+^ changes from (a). (c) The percentage of progressive motility in spermatozoa derived from donors determined by CASA before or after treatment with exosomes isolated from normozoospermic or asthenozoospermic patients. HSA stimulation served as a positive control. (d) The ability of sperm to penetrate into artificial viscous media was evaluated by above mentioned methods. Data were presented as mean ± SME (n = 5-10). *P < 0.05, **P < 0.01, ***P < 0.001 versus no treatment group (HTF). ##P < 0.01 versus normozoospermic exosome-treated group.

## Discussion

In this study, we for the first time demonstrated that the SPE-evoked [Ca^2+^]i resulted from extracellular Ca^2+^ influx, because exosomes had no effect on sperm [Ca^2+^]i in Ca^2+^-free medium. The exosome-induced elevation of [Ca^2+^]i was controlled by CatSper activation, as a CatSper inhibitor significantly impaired the exosome-activated Ca^2+^ response, and there were no [Ca^2+^]i and CatSper current responses using exosome-stimulated CatSper deficient sperm. In addition, exosomes and their cargos promoted sperm motility, especially hyperactivation.

This study used various methods, such as the Flow NanoAnalyzer, western blotting and TEM, to systematically identify and characterize exosomes isolated from human seminal plasma. Our data showed that we obtained exosomes with vesicle structural characteristics similar to those described in a previous study [35]. The exosome diameters (94.9 ± 21.8 nm) were consistent with past reports [6, 18]. However, the ratio of CD63 positive exosomes was approximately 18.5% in our samples, which may reflect a more complex composition and unique characteristics in human seminal plasma exosomes than those derived from other body fluids, and the present of some non-classical CD63 negative exosomal extracellular vesicles [35–37]. For examples, seminal fluid from some species, such as the chicken, lacks CD9- and CD44-bearing extracellular vesicles [38].

It was previously reported that exosomes contributed to elevated Ca^2+^ concentrations in sperm via the release of calcium contained in prostasomes during fusion with sperm, and it was proposed that a novel mechanism influenced sperm [Ca^2+^]i independently of ion-exchange systems and ATP-dependent pumps [31]. However, our results show that the exosome-induced sperm [Ca^2+^]i response was still determined by ion-exchange systems (CatSper). Although exosomes increased sperm [Ca^2+^]i in work by Palmerini et al., the elevated [Ca^2+^]i may have derived from the exosomes themselves, as we showed calcium contained in exosomes could also be monitored by the fluorescent probe (Flou-4 AM) (Figure S4). Palmerini et al. detected [Ca^2+^]i after the exosomes fused to sperm [31], and under these conditions, the detection of a fluorescence signal may directly result from some exosomes retained or integrated onto sperm. Our results demonstrated an ion channel-controlled mechanism involving CatSper-mediated exosome-induced [Ca^2+^]i signal via extracellular Ca^2+^ influx. The fusion or stimulation of sperm with exosomes and the transfer of Ca^2+^-related cargos to the sperm surface did not cause the [Ca^2+^]i response if CatSper didn’t work.

Exosomes reportedly contained proteins, nucleic acid, lipids and some small molecule compounds, such as prostaglandins [32, 39, 40]. Our results showed that triton X-100 and ultrasonic-treated exosomes conferred a stronger stimulatory effect on the [Ca^2+^]i signal response compared with intact exosomes. The precipitates isolated from triton X-100-treated exosomes had no effect on sperm [Ca^2+^]i. Furthermore, we found that exosome protein components contributed to the increased [Ca^2+^]i, because enhancement of [Ca^2+^]i was partially impaired using Pro K- or heat-treated exosome supernatants compared with untreated exosome supernatants. We speculated that exosome protein and non-protein components exerted Ca^2+^ signal activation in human sperm. Our study also found that intact exosomes contained nanomole levels of PGE1 and PGE2, consistent with other reports [32]. Moreover, PGE1 and PGE2 induced the sperm [Ca^2+^]i signal in our study (Figure S2(b)–(e)), consistent with previous research [33]. The detected concentrations of PGE1 and PGE2 were lower than the concentrations required for an effect, so we propose that exosomes may carry PGE1, PGE2 and other prostaglandins, as well as other non-protein cargos, that combine to induce sperm [Ca^2+^]i. Furthermore, we found triton X-100-treated exosomes had no detectable level of PGE1, but had higher levels PGE2 (Figure S2(a)), suggesting that exosomes maintained the stability and physiological activity of cargos, such as PGE1. It has been reported that the use of liposomes to deliver PGEs slowed the release rate, increased the therapeutic effect and diminished adverse reactions [41, 42]. The effects of PGE1 on human sperm function may involve an open CatSper channel, because transient intracellular Ca^2+^ signals induced by PGE1 were fully inhibited by CatSper blockers [43]. Interestingly, the mechanism of PGE1-stimulated CatSper activation may involve a direct effect and differ from a progesterone receptor-dependent pathway [44]. Here, we found that exosomes still augmented a [Ca^2+^]i signal after the progesterone induced-calcium signal reached a maximum threshold (Figure S5), suggesting that exosomes exerted a distinct and/or complex activation pathway that in part differed from the progesterone-induced open CatSper channel. These findings may be attributed to the combined effect of multiple cargos in exosomes.

Considering the CatSper-mediated hyperactivation of sperm [34], which could be triggered by exosomes, we speculated that exosomes may enhance sperm motility parameters, especially hyperactivation. Our results demonstrated that exosomes facilitated sperm hyperactivation and augmented sperm penetration into a methylcellulose medium, suggesting that exosomes may enhance the ability of sperm to pass through the viscous female reproductive tract. Furthermore, our data indicated that physiological concentrations of exosomes promoted sperm total and progressive motility, reflecting physiological conditions at the early stage of ejaculation, confirming previous reports showing exosomes promoted sperm motility in an artificial buffer [15, 45]. A recent study reported that exosomes derived from normozoospermic but not from asthenozoospermic individuals improved spermatozoa motility and triggered capacitation, suggesting that semen quality was affected by male tract exosomes [18]. In this study, the authors proposed that exosomes carried proteins, such as CRISP1, involved in sperm motility regulation. Other exosome proteins, CD38 and cSrc kinases, also were thought to confer sperm hypermotility [46, 47]. In this study, we confirmed that protein components in exosomes regulate sperm motility, especially hyperactivation, because the hyperactivated sperm count and the sperm’s ability to pass through a viscous medium were impaired after exosome proteins were eliminated by Pro K or heat treatment.

Finally, we found that exosomes improved the motility of sperm from patients with asthenospermia, as determined by the elevated progressive motility and ability to penetrate viscous medium. This finding was consistent with previous work indicating that sperm motility can benefit from exosomes [28]. Nevertheless, the physiological significances of exosomes for sperm function regulation remain an intriguing and challenging topic. Exosomes with other seminal plasma and female reproductive tract components may together modify sperm function required for fertilization. We also found that that exosomes isolated from severe asthenozoospermic samples (PR < 15%) have a markedly weaker effect on [Ca^2+^]i activation than normal exosomes. This result, combined with the recent report, suggests the dysfunction and dyssecretosis of exosomes may contribute to low sperm motility and infertility [18]. Calcium regulation is crucial for sperm motility. Spermatozoa from some asthenozoospermic patients exhibited a reduced responsiveness to progesterone [48]. Hence, we propose that high quality endogenous or engineering-modified exosomes may be developed as a therapeutic agent to treat the lowered sperm motility caused by calcium signal response disorders.

In conclusion, SPE activated CatSper to induce [Ca^2+^]i in human sperm via extracellular Ca^2+^ influx, which contributed to exosome-induced hyperactivation. Exosomes provide cargos that allow sperm to mobilize and manage a CatSper-regulated Ca^2+^ signaling mechanism in sperm. The improvement of asthenozoospermic sperm motility exerted by exosomes provides a new potential clinical opportunity. This study provides insight into the role and mechanism of human SPE in sperm function, and reveals a novel CatSper regulator for further examination in cases of sperm dysfunction and male infertility caused by [Ca^2+^]i dysregulation.

## Supplementary data

Figure S1. Exosome Cargos induced-[Ca^2+^]i increase via extracellular Ca^2+^ influx.

Figure S2. Contents of PGE1 and PGE2 in exosomes and effect of them on sperm [Ca^2+^]i.

Figure S3. Effect of exosomes on sperm motility.

Figure S4. Calcium contained in exosomes could be monitored by Flou-4 AM.

Figure S5. Synergistic effect of progesterone with exosomes on sperm [Ca^2+^]i response.

## Acknowledgments

The authors thank all the participants who took part in this study, as well as all medical staff who helped with human semen collection in Nanchang Reproductive Hospital and Jiangxi Maternal and Child Health Hospital. We thank Charles Allan, PhD, from Liwen Bianji, Edanz Group China (www.liwenbianji.cn/ac), for editing the English text of a draft of this manuscript.

## Author Contributions

X.H.Z., D.D.S. and X.N.Z. designed the study and wrote the manuscript with feedback from the other authors. H. K. performed sperm patch-clamp recordings. D.D. S. and W.W. Z. participated in single cell calcium imaging, the penetration of artificial viscous media assay and CASA, hyperactivation, WB and the detailed data analysis. H.Y.C. and H. K. acquired and processed all patient samples. X.H.Z. and X.N.Z. were responsible for primary study oversight, data analysis, interpretation and manuscript revision. All authors read and approved the final version of the manuscript for submission.

---

Figure S1. The exosome cargos induced-[Ca^2+^]i increase in sperm depended on extracellular Ca^2+^ influx. Sperm [Ca^2+^]i were recorded according to above mentioned methods. The data were cumulative of at least three independent experiments. *P < 0.05, **P < 0.01, ***P < 0.001 versus HS. ###P<0.001 versus exosomes.

Figure S2. Contents of PGE1 and PGE2 in exosomes and effect of them on sperm [Ca^2+^]i. (a) Contents of PGE1 and PGE2 were examined by Elisa Kits. (b)-(e) Sperm [Ca^2+^]i were recorded after stimulated to PGE1 or PGE2 according to above mentioned methods. The data were cumulative of at least three independent experiments. *P < 0.05, **P < 0.01, ***P < 0.001 versus HS.

Figure S3. Effect of exosomes on sperm motility. Human sperm were incubated in seminal plasma or exosomes-free seminal plasma at 37 °C and 5% CO_2_ incubator for different time. The motion parameters,(a) total motility, (b) PR, (c) VCL, (d) VSL, (e) VAP, (f) LIN, were analyzed by CASA. A minimum of 200 sperms were counted for each assay. The data were cumulative of at least three independent experiments. *P < 0.05, **P < 0.01, ***P < 0.001 versus seminal plasma.

Figure S4. Calcium contained in exosomes. The fluorescence intensity of Ca^2+^ in exosomes was detected by Multimode Plate Reader with Fluo 4 (2 μM) staining.

Figure S5. Synergistic effect of progesterone with exosomes on sperm [Ca^2+^]i response. Sperm [Ca^2+^]i were recorded according to above mentioned methods. The data were cumulative of at least three independent experiments.

