## Supplementary figures and images for "Seminal plasma exosomes evoke calcium signals via the CatSper channel to regulate human sperm function"

### Figure S1

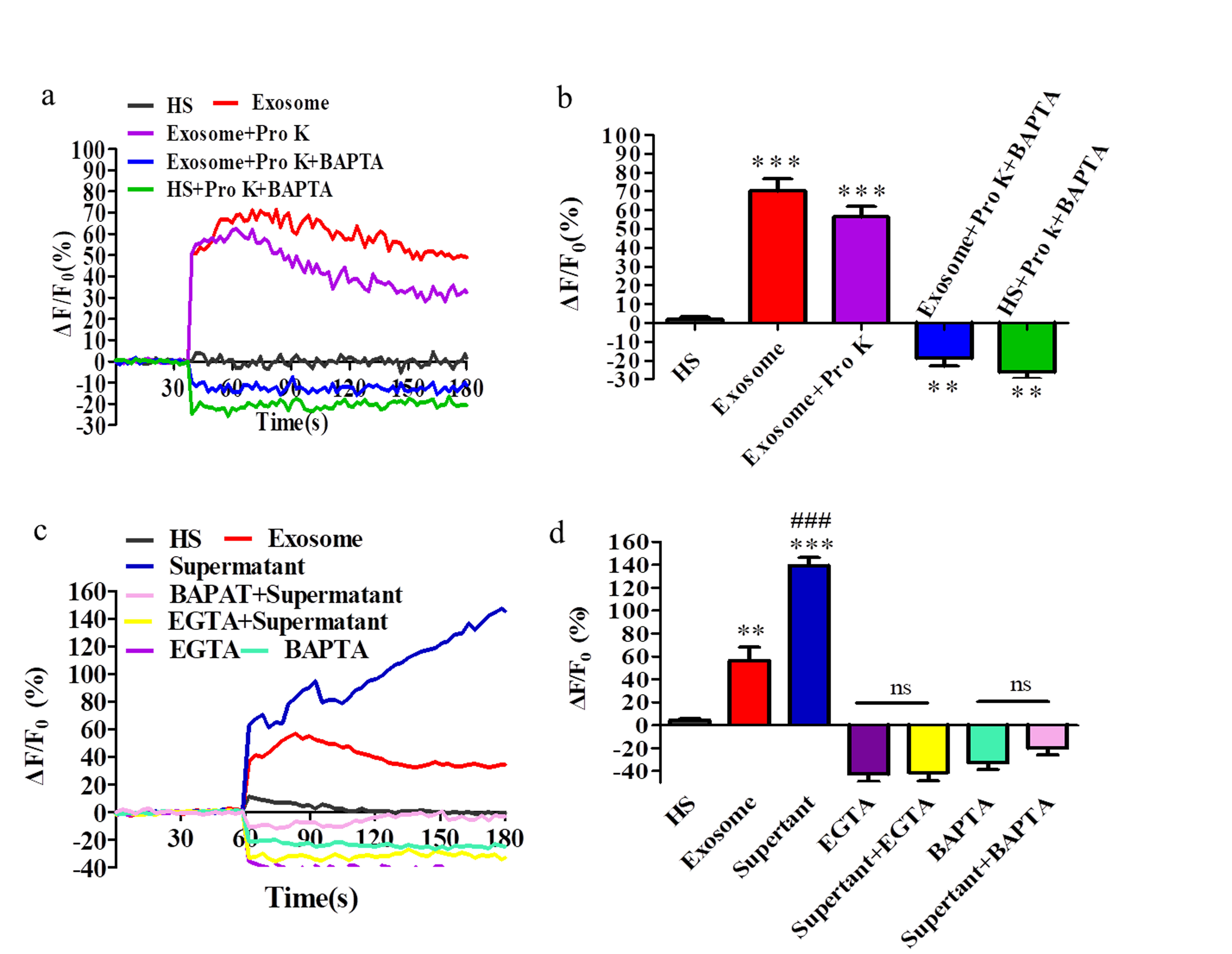

### Figure S2

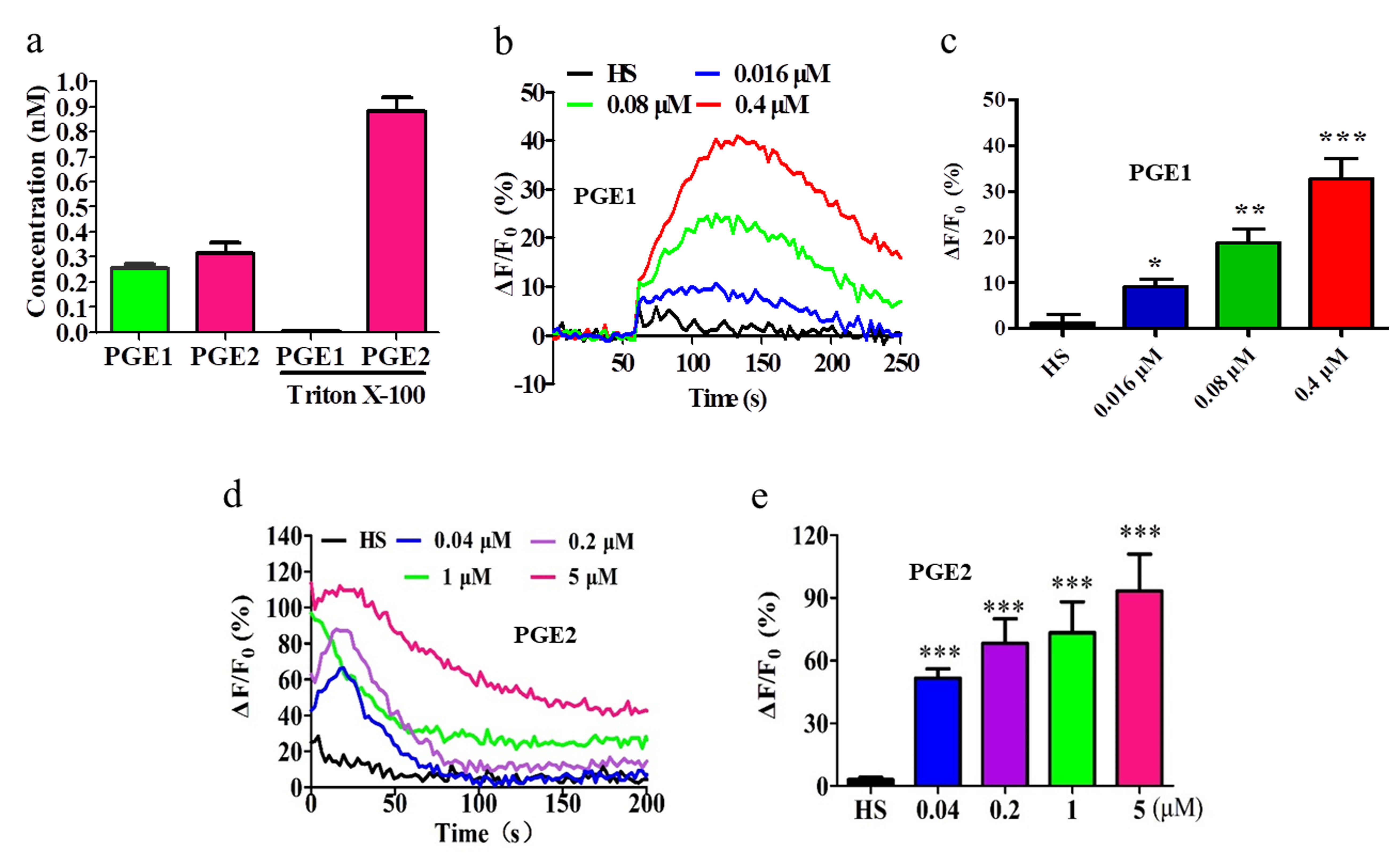

### Figure S3

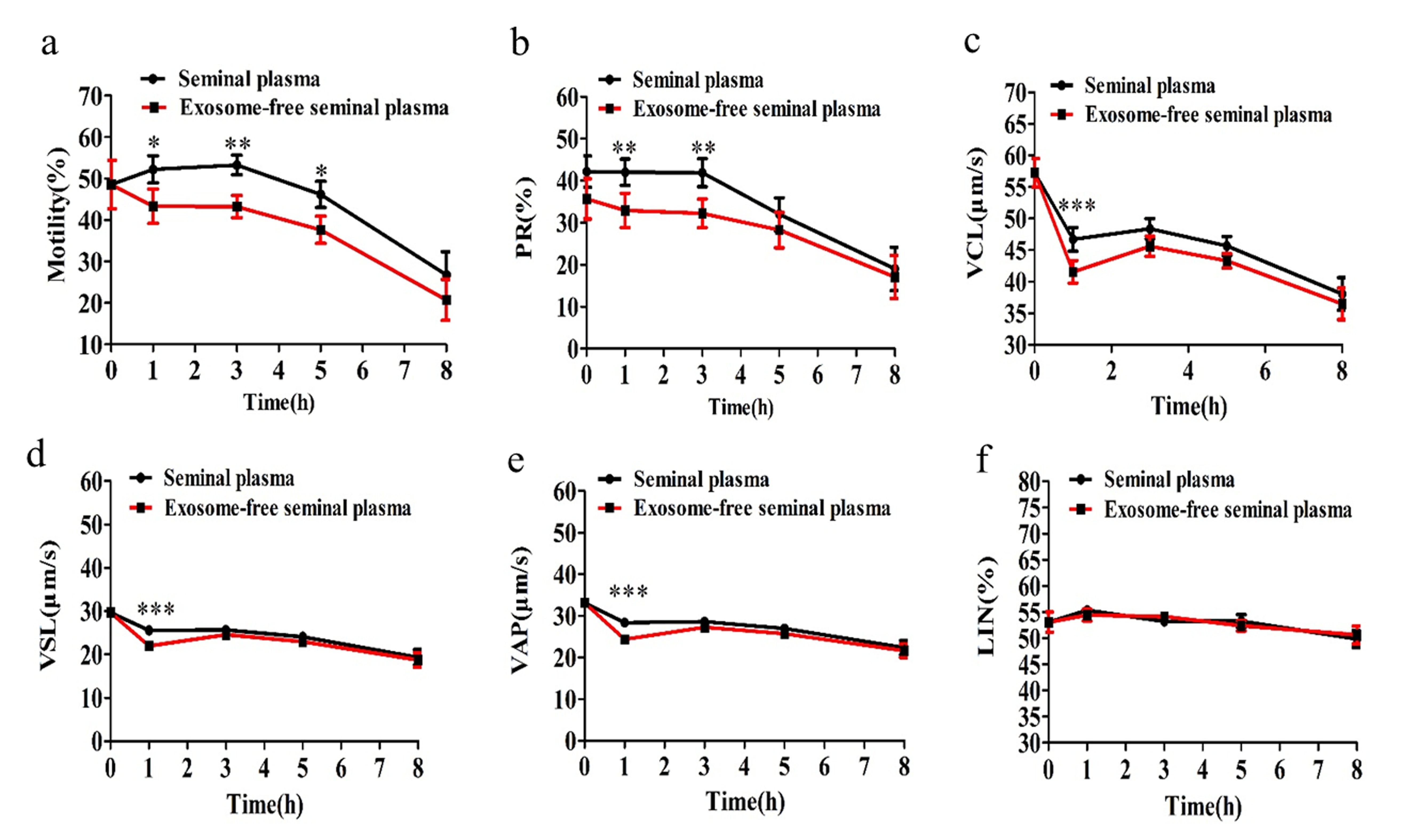

### Figure S4

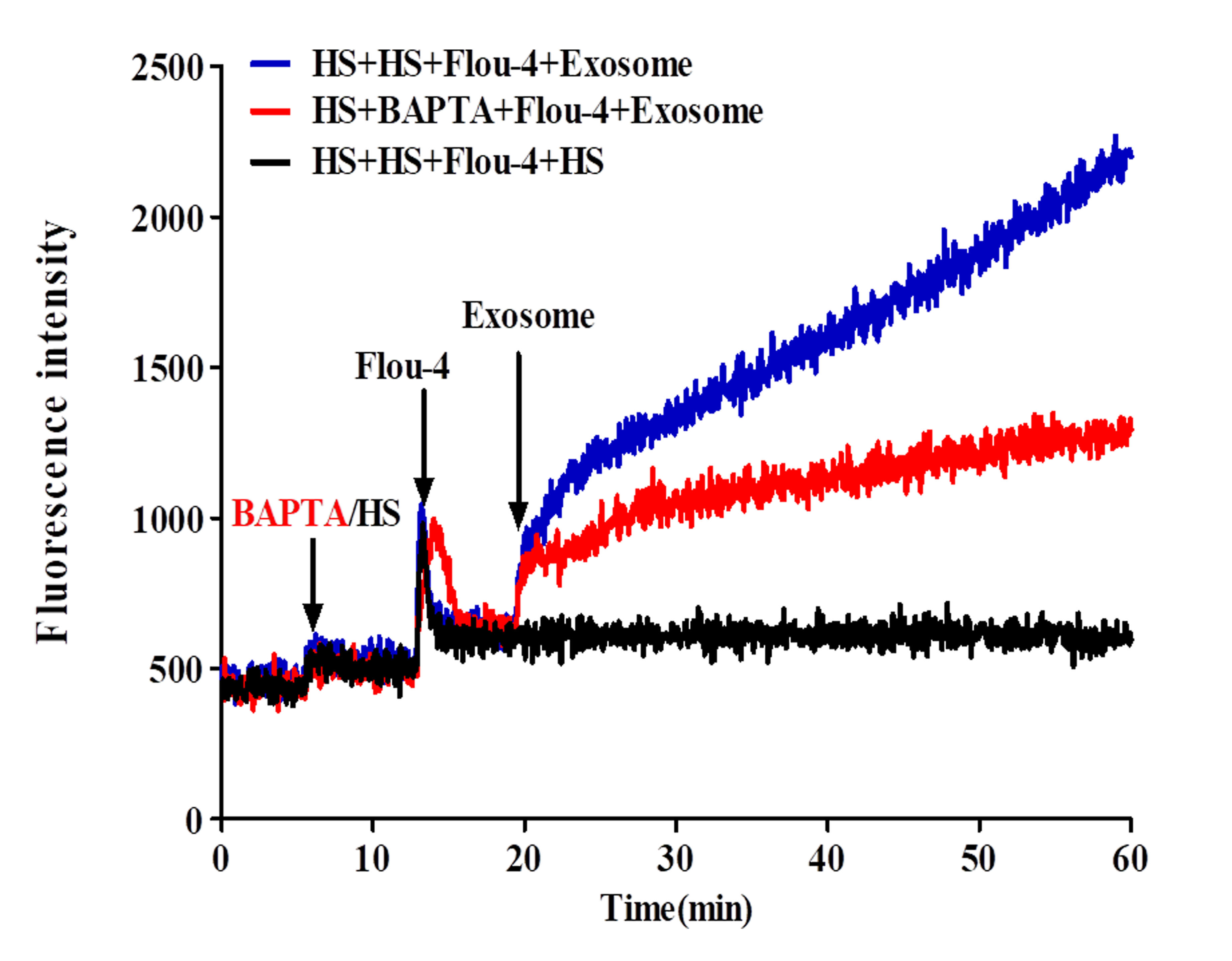

### Figure S5

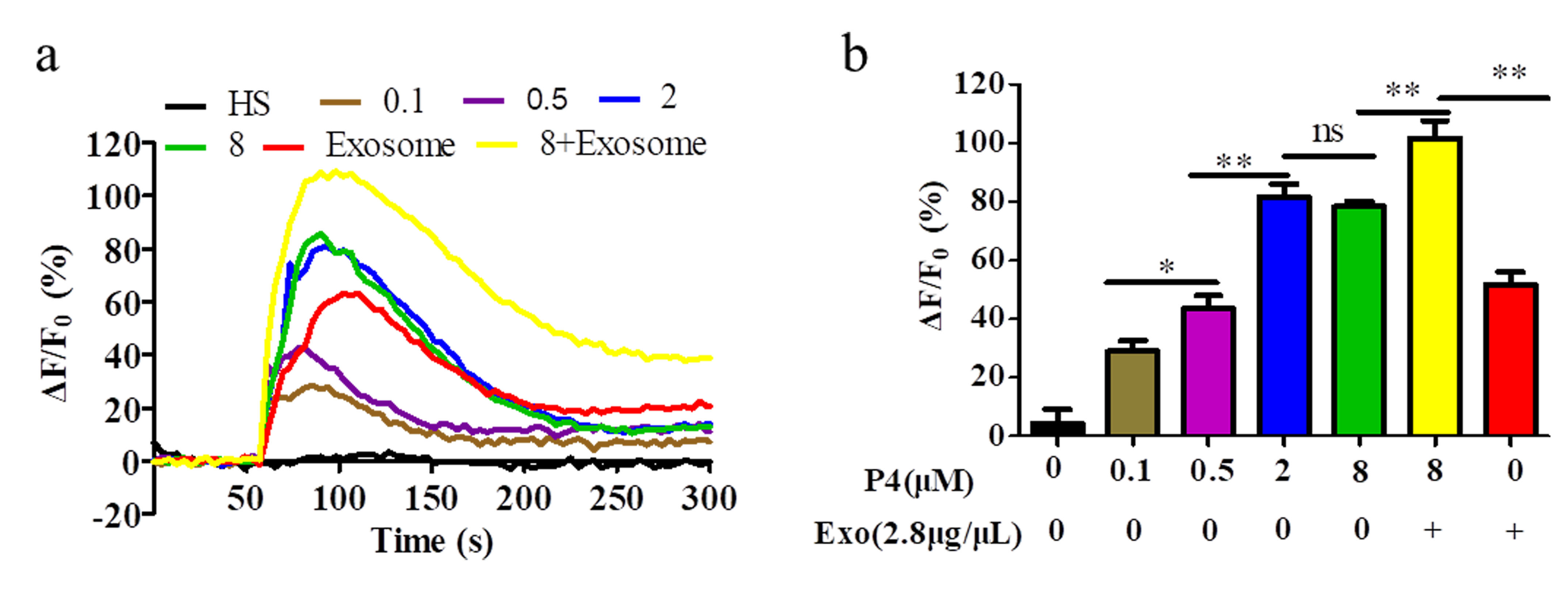
